# Active site center redesign increases protein stability preserving catalysis

**DOI:** 10.1101/2022.01.31.478513

**Authors:** Maria Luisa Romero-Romero, Hector Garcia-Seisdedos, Beatriz Ibarra-Molero

## Abstract

The stabilization of natural proteins is a long-standing desired goal in protein engineering. Optimizing the hydrophobicity of the protein core often results in extensive stability enhancements. However, the presence of totally or partially buried catalytic charged residues, essential for protein function, has limited the applicability of this strategy. Here, focusing on the thioredoxin, we aimed to augment protein stability by removing buried charged residues in the active site without loss of catalytic activity. To this end, we performed a charged-to-hydrophobic substitution of a buried and functional group, resulting in a significant stability increase yet abolishing catalytic activity. Then, to simulate the catalytic role of the buried ionizable group, we designed a combinatorial library of variants targeting a set of seven surface residues adjacent to the active site. Notably, more than 50% of the library variants restored, to some extent, the catalytic activity. The combination of experimental study of 2% of the library with the prediction of the whole mutational space by Partial Least-squares regression revealed that a single point mutation at the protein surface is sufficient to fully restore the catalytic activity without thermostability cost. As a result, we engineered one of the highest thermal stability reported for a protein with a natural occurring fold (138 °C). Further, our hyperstable variant preserves the catalytic activity both *in vitro* and *in vivo*.

**SIGNIFICANCE:** The major driving force of protein folding is the hydrophobic effect, and increasing the protein core hydrophobicity essentially increases protein stability. Active sites often contain buried ionizable groups, which can be essential for function but dramatically reduce protein stability. Thus, increasing the protein core hydrophobicity cannot be applied to enzyme active sites without a functional cost. We propose a method to enhance protein stability by overcoming this obstacle. We show that catalytic properties of buried charges can be mimicked with surface mutations, thus paving the way to unlock the optimization of the hydrophobic core to stabilize enzymes.

## INTRODUCTION

Most proteins exhibit nearly balanced free energy profiles for folded and unfolded states^1^ which hampers their biotechnological applications. Furthermore, as most mutations are destabilizing^2^, marginal stability becomes a significant bottleneck for the laboratory evolution^3,4^ and computational design of proteins^5,6^. In consequence, many efforts have been directed to increase protein stability^1,7–9^. Notably, *de novo* design has been revealed as one of the most successful strategies to achieve protein hyperstability, partly attributed to the well-packed and exclusively hydrophobic cores of its designs^10^. Indeed, optimizing natural proteins’ core to maximize its buried hydrophobic surface area often results in extensive stability enhancements11–16. However, it is frequently impossible to fully implement such a strategy in natural proteins, as buried ionizable residues are often part of active sites and, thus, linked to function17–19. Therefore, when applying this stability strategy to natural proteins, functional buried ionizable residues typically remain untouched.

This trade-off between stability and function is exemplified by the electron transfer protein thioredoxin. The thioredoxin^29^ (Trx) is a small and well-characterised protein with an active site center partially exposed^30^. It comprises a tryptophan followed by two vicinal cysteines spaced by a glycine-proline segment (Trp-Cys_32_-Gly-Pro-Cys_35_), enumerated according to the sequence of the *E.coli-Trx*, and a conserved and buried aspartic residue (Asp_26_)29–31. Mutating the buried Asp_26_ to a hydrophobic residue dramatically drops the catalytic activity31,37 yet increases the protein stability13,29.

We aim to increase protein thermostability by replacing a buried and functional charged group with a hydrophobic residue without stability cost. It has been previously shown how the rational redesign of the surface charge distribution can modulate the p*K*_a_ and the ionization state of a buried residue^28^. In this work, we go a step further and aim to simulate a buried ionizable group’s functional role, comprised in the active-site center of a protein, by redesigning the surface charge distribution adjacent to the active site. Such a strategy may allow maximizing the buried hydrophobic area of proteins while preserving their catalytic capabilities.

Here, we optimized the thioredoxin’s protein core by mutating its buried and functional Asp_26_ to Ile. Then, based on structural criteria, we selected a set of solvent-exposed positions that might balance the absence of the buried aspartic residue. After that, we constructed a combinatorial library of thioredoxin variants where the previously selected positions were mutated to lysine, glutamate, or kept the Wild-type residue. We tested the thermostability and catalytic activity of 39 variants accounting for 1.8% of the total library of 2187 variants. We used partial least-squares (PLS) reconstruction to predict the whole mutational space of the library. Lastly, we selected those variants with predicted higher redox activity and experimentally validated their thermostability and activity *in vitro* and *in vivo.*

Notably, our results show that about half of the library’s variants restored the catalytic activity of the thioredoxin to some degree. Strikingly, introducing a sole lysine on the protein surface was enough to completely restore the activity of the conserved and buried aspartic residue *in vitro.* This redesigned active site variant was functional *in vivo* and displayed one of the highest thermal stability ever reported^32^. These findings support that neutral-to-charged mutations of solvent-exposed residues can mimic the select physicochemical properties of catalytic buried ionizable groups. This potential might be harnessed to stabilize natural proteins by substituting functional yet destabilizing buried charged groups without compromising their catalytic activities.

## RESULTS

### Surface mutations simulate the catalytic role of a buried charged residue in thioredoxin *in vitro*

We want to increase protein stability maximizing the protein core’s hydrophobicity. An established approach consists in substituting buried charged residues with hydrophobics^11–16^. However, as buried charged groups are often engaged in functional roles, this strategy risks augmenting stability at the expense of function. Therefore, first, we aim to fashion the functional role of a buried charged residue by surface mutations directed to redesign the charge surface distribution nearby. This strategy profits from the plasticity of the protein’s surface, which allows rearranging the protein, minimizing the stability effect of the mutations^27^.

To meet this goal, we focused on thioredoxin. Thioredoxins are proteins present in all organisms that catalyze disulphide bond reductions^33^. Their canonical active site center hosts the segment Trp-Cys_32_-Gly-Pro-Cys_35_, where the two cysteine’s side-chain atoms are mostly buried. Close to the disulfide and part of the active-site center, a buried carboxylic residue (Asp_26_) is utterly conserved among thioredoxins (**Fig 1a**). The reaction catalyzed by Trx is a bimolecular nucleophilic substitution that involves shuttling two electrons from the Trx to the substrate. Firstly, the hydrophobic active site residues interact with the substrate protein. Then, in the hydrophobic environment of the complex, Cys_32_ nucleophilically attacks the substrate’s disulfide bond, resulting in an intermediate disulfide bond between Trx and the substrate. Finally, the deprotonated Cys^35^ nucleophilically attacks the mixed disulfide bond, resulting in the oxidation of Trx and the reduction of the substrate^34^ (**Fig 1b**). The buried Asp_26_ deprotonates and activates the Cys_35_^35–37^, guiding the reaction’s molecular mechanism and removing the charge of residue 26 results in a dramatic decrease in the redox capacity of Trx^31^. To test whether we can mimic the contribution to the redox activity of Asp_26_, we used as the library starting point a modified version of an ancestral Trx from the last universal common ancestor of the Cyanobacterial, Deinococcus, and Thermus groups^38^, called from now on trx0, that has a melting temperature (T_m_) of 128°C29. Trx0 will likely accept several destabilizing mutations, allowing to explore a more expansive mutational space and, therefore, increasing the chances to find an alternative active site center to that found in all-natural thioredoxins.

**Figure 1.**
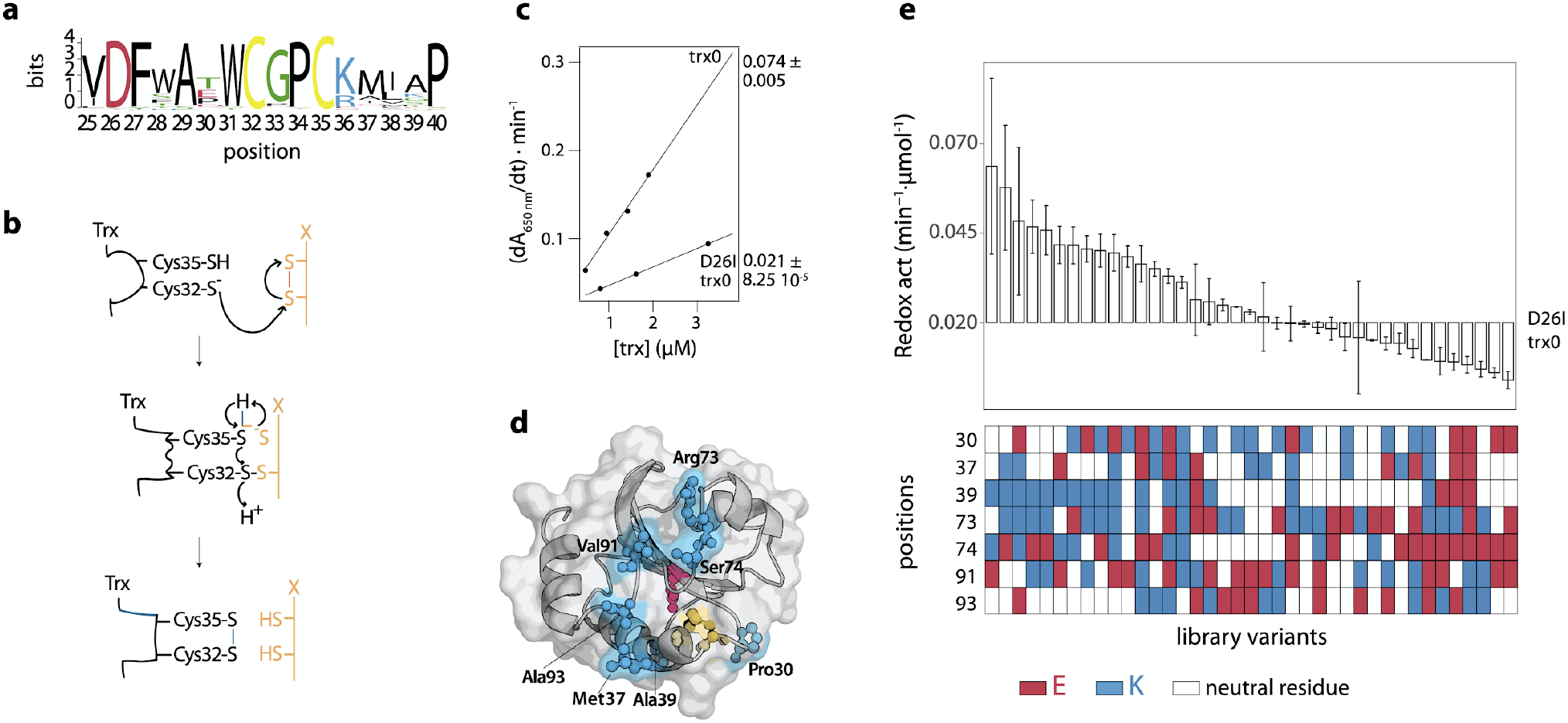
Redesign of the conserved active site of the thioredoxin. **a)** Sequence logo showing the conservation of the thioredoxin active site. The logo was inferred from an alignment of 120 thioredoxin sequences which shared at least 30% of identity with E. *coli* Trx **b)** Molecular mechanism of the redox activity of the thioredoxin. Reduced thioredoxin binds, through its hydrophobic active site area, to the substrate. Then, the thiolate of Cys_32_ attacks the substrate and forms a transient mixed disulfide. After that, the thiolate of Cys_32_ attacks this mixed disulfide, generating the oxidized thioredoxin and the reduced substrate protein^34^. **c)**. The redox activity of the thioredoxins was determined testing their capacity to reduce insulin disulphide bridges. The activity values are determined from the slope of the linear relationship between the maximum rate of the insulin reduction curves versus thioredoxin concentration. The errors shown correspond to the standard deviation associated with the slope. The mutation D26I significantly drops the redox activity of trx0. **d)** Crystallographic structure of the ancestral thioredoxin from the last common ancestor of the cyanobacterial, *Deinococcus* and *Thermus* groups (LPBCA, pdb code: 2YJ7)^39^. The catalytic, partially buried cysteines are shown in yellow, and the catalytic buried aspartic in red. Those solvent-exposed residues selected to modulate the functional role of the buried Asp_26_ are shown in cian. **e)** Experimental redox activity of the 39 randomly selected variants of the library. The activity values are determined from the slope of the linear relationship between the maximum rate of the insulin reduction curves versus thioredoxin concentration. The error is the standard deviation associated with the slope. As a color-code, below is shown the type of residues found at the seven positions targeted in the library. White when there is a neutral residue, blue when there is glutamine, and red when there is a lysine.

Firstly, we measured the effect of the D26I substitution on trx0 activity. The D26I substitution reduced the redox capacity by 72% (**Fig 1c**), consistent with the previously described functional role of the buried aspartic residue in *E.coli* Trx31,37 and the absolute conservation of the thioredoxin structure over evolution^39^. Then, we measured the thermostability of D26I-trx0. The D26 substitution increases by 10°C the protein stability, scaling the T_m_ up to 138°C, in agreement with the previously reported stability effect of the D26I mutation in *E.coli* Trx13. Thus, D26I-trx0 is a hyperstable protein with a residual capacity to catalyze the reduction of disulphide bonds.

We selected a set of residues that, upon mutation, might simulate the buried aspartic’s functional role based on structural criteria: 1- We selected those residues close to the active site center. For this purpose, we picked all residues in a radius of 11 Å centered in the sulfur atom of the catalytic Cys_32_. 2- Among those residues, we discarded the buried ones, *i.e.*, we only considered residues with an accessible surface area (ASA) above 0.3. 3- Lastly, we excluded exposed residues directly involved in the redox activity, such as Trp_31_, Asp_61_, Pro_76_, and Lys_57_^40^, and those located adjacent to the sulfur atom of the Cys_32_ – closer than 7 Å. With these criteria, seven positions were selected as promising candidates to change the surface charge distribution and simulate the catalytic role of the buried Asp_26_: Pro_30_, Met_37_, Ala_39_, Arg_73_, Ser_74_, Val_91_, and Ala_93_ (**Fig 1d** and **Table 1**). We then designed a combinatorial library of variants where each position could be occupied by a neutral, a positive, or a negative charged residue. Positions 30, 37, 39, 74, 91, and 93 could be hosted by their wild type neutral residue or mutated either to lysine or glutamic acid. However, the wild type residue found in position 73 is already positively charged; therefore, in the library, this position could be occupied by its wild type residue or mutated either to glutamic acid or to methionine (since methionine is often found in this position in an alignment of thioredoxins).

**Table 1.** Selected solvent-exposed positions to simulate the functional properties of the buried Asp_26_. In the first column is annotated the amino acids found in the position shown in the second column. In the third column it is written the distance from the sulfur atom of the catalytic Cys32 to the closest atom of the aminoacid at each position, and in the fourth column, the accessible surface area.

| residue | position | $d_{Cys_{32}} (\text{\AA})$ | ASA ( $\text{\AA}^2$ ) |
| --- | --- | --- | --- |
| proline | 30 | 7.0 | 0.58 |
| methionine | 37 | 9.4 | 0.73 |
| alanine | 39 | 9.5 | 0.33 |
| arginine | 73 | 10.1 | 0.93 |
| serine | 74 | 7.5 | 0.58 |
| valine | 91 | 10.2 | 0.56 |
| alanine | 93 | 8.4 | 0.33 |

We constructed the combinatorial library and transformed it into E. *coli* cells. We randomly selected 104 colonies, from which 43 expressed soluble thioredoxin variants. Sequencing of the soluble variants revealed 39 unique mutants. We purified these mutants and studied their redox activity *in vitro* using insulin as a substrate^41^. Remarkably, nearly half of them showed enhanced redox activity compared to D26I-trx0. Indeed, some mutants displayed up to 3-fold redox activity than D26I-trx0, almost fully compensating for the lack of the buried Asp_26_ (**Fig 1e, Fig S1**). The results showed that redesigning the solvent-exposed charge distribution adjacent to the active site might allow the simulation of the catalytic properties of buried charged residues.

### Partial-least-square allows the phenotypic prediction of the entire mutational space

Complete experimental characterization of the whole library is infeasible. We still did not determine most variants’ activity (more than 98% of the library). Hence, we applied a computational method that allows the whole library’s phenotypic prediction from the 1.8% experimentally studied. We fit the experimental data to the equation:

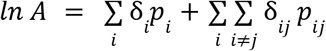

Where A is the redox activity; *δ_i_* is a vector of values (1, 0, 0) when the position *i* is occupied by a neutral residue, (0, 1, 0) when positively charged residue occupies it, and (0, 0, 1) when a negatively charged residue occupies it; *p_i_* is the effect of the residue in the position *i* on the redox activity; δ_*ij*_ = δ_*i*_·δ_*j*_, takes different values depending on the type of residue found in position *i* and *j* (**Table S1**) and *p_ij_* describes the coupling effect between residues at positions *i* and *j* in the redox activity. Hence, the first term of the equation describes the effect of the individual mutations on the activity and the second, the coupling effect of the mutations on the activity, considering a possible non-additive effect. The fitting requires 231 output parameters (21 *p_i_* parameters and 210 *p_ij_* parameters). At the same time, the number of experimental values to be fitted is only 39 (values of the redox activity of the screened library variants). We addressed the fitting using partial least-squares (PLS), an established algorithm to describe the relationship between protein sequence and function^42–44^. This approach has been used to predict the phenotypic outcome of the presence or absence of mutations42,43. Here, we further increase the mutational space to explore three different alternatives per position (a neutral, a positive, and a negatively charged amino acid) and consider the synergisms effect among mutations. PLS can predict a significant number of fitting parameters using very few experimental data because it first reduces the number of independent variables (*p_i_* and *p_ij_*) to fewer latent variables^45^. These latent variables are orthogonal combinations of the original independent variables that define most of their variance and explain the correlation between the independent and dependent variables (A, the redox activity). Thus, PLS uses the two data sets’ information, the dependent and independent variables, to define latent variables that maximize prediction capacity.

We bootstrapped the experimental data set to assess the prediction’s uncertainty, *i.e.*, we repeated the PLS fitting using 10 different data sets by resampling the 39 initial experimental values. Full-library reconstructions of such 10 replicas are shown in **Fig 2a**. The reconstruction displayed an unexpected scenario, and the redox activity might reach wild-type levels with very few mutations. We validated the reconstruction by selecting the best-predicted variants with one or two mutations (A39K, A39E, P30KA39K/S74K, A39K/K73M, and A39K/P30K) and experimentally examined their redox activity *in vitro* (**Fig 2c**). Remarkably, a single point mutation alanine-to-lysine in a solvent-exposed position (A39K) almost fully compensates for the functional properties of the buried Asp_26_, suggesting remarkable plasticity of the thioredoxin’s active site center. Further, the A39K mutation involves the accommodation of an ionizable residue in the protein surface, and thus, it likely implies a minor stability effect.

**Figure 2.**
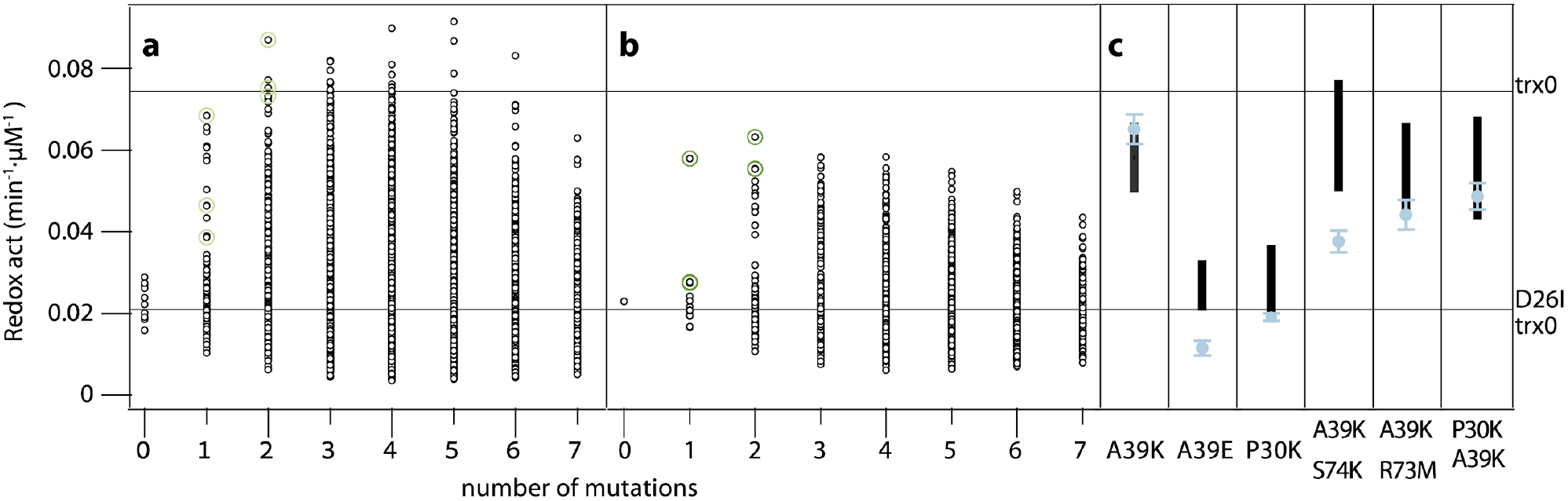
Partial least-squares reconstruction of the redox activity of all the library variants. **a)** Full-library reconstruction of the 10 data sets, obtained by resampling the 39 initial experimental values, are shown in black. In light green are highlighted the highest predictions for those variants with one or two mutations. **b)** In black, mean values of the predicted redox activity for each library variant. In green, mean values of the predicted redox activity for the best variants with one or two mutations. **c)** For each selected variant, in black is shown the range of predicted values and in blue the experimental value for the redox activity.

### Simultaneous enhancement of redox activity and stability in thioredoxin

We showed that surface mutations could reproduce the role of a buried charge on the redox activity of thioredoxin *in vitro*. However, the mutations’ stability effect and the interplay between stability and activity remain to be tested.

To understand the energetic landscape confined by the library variants, we studied their thermostability by differential scanning calorimetry (DSC) (**Fig S2**). Few variants displayed thermal stabilities equivalent to trx0. Remarkably, the A39K substitution, with similar catalytic activity to trx0, presented a T_m_ of 138 °C (**Fig 3a**). Therefore, we wondered whether the redesigned active-site center allows the multi-feature optimization of the enzyme, *i.e.*, whether simultaneously improving stability and activity is feasible.

**Figure 3.**
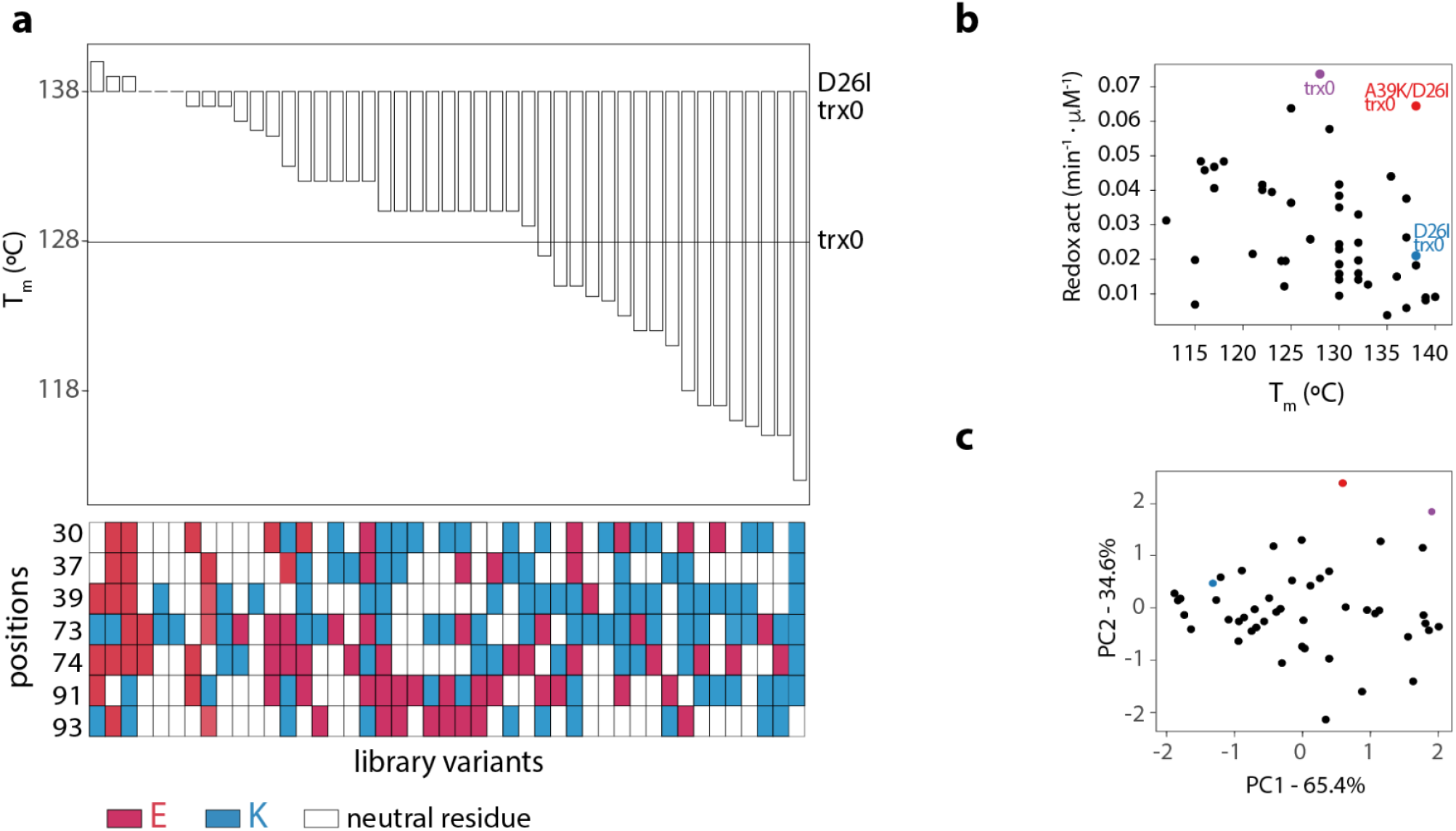
Redesigning the conserved active-site center allows the multi-feature optimization of the enzyme. **a)** Most of the variants decrease the thermostability of the library’s background – D26I-trx0. However, compared with trx0, half of the variants have improved stability. **b)** Plotting the redox activity versus the thermostability of each variant displays a spread out distribution of the experimental dataset. **c)** The PCA analysis displays two components with significant size. PC1 explained the 65% of the variance, whereas the PC2 explained the 35%.

To analyze whether stability and activity trade-off, we studied the stability-activity correlation among the experimentally studied variants. Plotting both properties shows, at a glance, a spread-out distribution of the dataset (**Fig 3b**). To quantify the degree of interplay between properties, we carried out a principal component analysis (PCA)46. PCA is a mathematical algorithm that reduces the dataset’s dimensionality, preserving as much statistical information as possible. It identifies directions, called principal components, along which the variation in the data is maximal. If two properties correlate, the dataset can be reduced to only one component. Hence, PCA displays a major component and a very minor one. On the contrary, if two properties are independent, two components are needed to explain the data variation. Then, the PCA analysis displays two components with significant size. Two components contributed significantly to the variance of the stability-activity dataset. PC1 explained 65% of the variance, whereas PC2 explained 35% (**Fig 3c**). Thus, the PCA analysis revealed a weak correlation and an efficient independent modulation of both properties. Overall, the modulation range achieved for the catalysis and stability was broad and, since both parameters weakly correlate, a simultaneous improvement of activity and stability seems possible. Indeed, the mutant A39K, whose redox activity is comparable to trx0 and higher than that of *E.coli* and human thioredoxins^38^, has a T_m_ of 138 °C.

Taken together, the results show that the small impact of surface mutations on protein stability permits improving the catalytic activity without compromising the stability.

### Synthetic thioredoxin functions *in vivo*

The mutation A39K in thioredoxin reproduces the catalytic role of the buried Asp_26_ *in vitro* and stabilizes the protein scaffold. We wondered whether our redesigned variant is active in a cellular context. In the cell, thioredoxin must catalyze the reduction of disulfide bridges of numerous protein substrates, and multiple factors such as crowding and viscosity are at play. To assess whether the redesigned thioredoxin (D26I/A39K-trx0) functions *in vivo*, we genome-integrated selected thioredoxin variants (trx0, D26I-trx0, and D26I/A39K-trx0) in a *Saccharomyces cerevisiae* strain lacking the TRX2 gene^47^. TRX2 encodes the protein thioredoxin II, which maintains thiol peroxidases in the reduced state. Consequently, the TRX2 deletion mutant (trx2Δ) is hypersensitive to oxidative stress^48^. Thus, if our thioredoxin variants are active *in vivo*, they should, to some extent, complement the function of trx2 under oxidative stress. As a positive control, we also genome-integrated the yeast TRX2. As thioredoxin is a moonlighting protein^49^ with multiple interaction partners50–52, a possible increase in fitness could be partially attributed to specific interactions or functionalities independent from the active site. To minimize this effect we conduct the experiments in yeast, an organism that is evolutively distant from trx0, which is derived from an ancestral thioredoxin corresponding to an internal bacterial node^29^. Indeed, while trx0 and *E. coli’*s thioredoxin share an identity of 58.3%, the identity shared by trx0 and yeast’s trx2 is only 38.3%.

We studied the growth of these mutants in normal growing conditions and under oxidative stress. The lag time of the trx2Δ strain increased 3.3-fold and 6.3-fold in the presence of 3 mM H_2_O_2_ and 4 mM H_2_O_2_ respectively (**Fig 4a**, **S3, and Table S2**). We observed a similar extension of the lag phase under H_2_O_2_ exposure for the D26I-trx0 variant (3.0-fold increase at 3 mM H_2_O_2_ and 6.0-fold at 4 mM H_2_O_2_), yet the lag phase was reduced for trx0 (2.7-fold and 5.2-fold increase at 3 and 4 mM H_2_O_2_) and even more for the redesigned thioredoxin D26I/A39K-trx0 (2.6-fold and 5.0-fold at 3 and 4mM H_2_O_2_) (**Fig 4a and Table S2**). Notably, the expression of D26I/A39K-trx0 decreased the lag-phase in 3.2h at 3 mM H_2_O_2_ and in 9.1h at 4 mM H_2_O_2_ with respect to D26I-trx0. While D26I/A39K-trx0 was not able to suppress the peroxide sensitivity to the same degree than trx2 (1.8-fold and 2.5-fold increase in the lag-phase at 3 and 4mM H_2_O compared with the lag-phase at 0 mM H_2_O), it displayed similar positive complementation as trx0 and significantly improved compare with D26I-trx0. The trx0 is an ancestral thioredoxin variant that was supposed to exist in the Precambrian era29,38. The coevolution of the thioredoxin with the organism’s protein network is fundamental to optimizing its performance. This is reflected in the superior capacity of endogenous thioredoxin (trx2) compared with D26I/A39K-trx0. However, D26I/A39K-trx0 significantly raises the redox activity *in vivo* with respect to trx2Δ and D26I-trx0.

**Figure 4.**
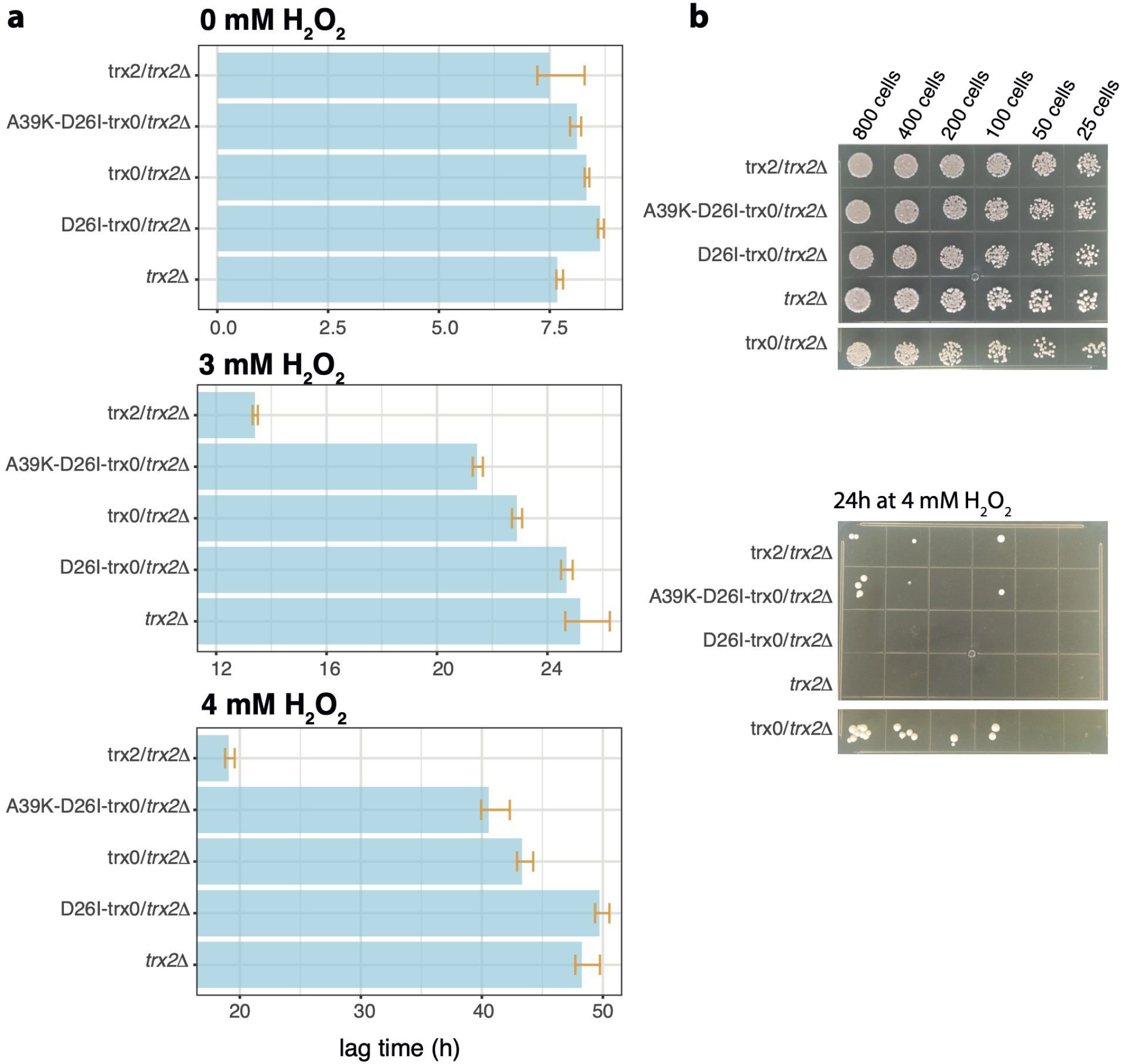
The redesigned thioredoxin, D26I/A39K-trx0, can catalyze the reduction of disulfide bonds *in vivo*. **a)** *In vivo* complementation assays in yeast displayed the following scenario. The growth of trx2Δ, presents a long lag-phase when stressing with H_2_O_2_. This lag-phase is maintained when overexpressing D26I-trx0. However, the lag-phase at the highest H_2_O_2_ concentration is shortened in 5h, 8h and 29h with the overexpression of trx0, A39K/D26I-trx0 and trx2, respectively. **b)** Plate-based spot assay to visualize yeast cells’ survival after treatment with 4 H_2_O_2_ for 24 h at 30°C. *trx2Δ* and *D26I/trx2Δ* could not survive the peroxided stress. Overexpression of trx0, D26I/A39K and trx2 allows surviving the peroxide treatment. In the upper panel is shown the number of colonies before the treatment with 4 H_2_O_2_ and below, after 24h at 30°C with YPD supplemented with 4 H_2_O_2_.

The growth of the trx2Δ yeast constructs presented a lag-phase phenotype in the presence of H_2_O_2_ while no phenotype in the exponential or stationary phase (**Fig S3**). However, stress response assays can be highly variable depending on cell growth conditions^53^, as observed by the elongated error bars, especially at the exponential and stationary phases (**Fig S3**). Thus, we performed a second assay for oxidative stress induced by hydrogen peroxide treatment, optimized to obtain highly-reproducible results^54^. This plate-based spot assay allows the visualization of yeast cells’ survival after treatment with 4 mM H_2_O_2_ for 24 h at 30°C. The results showed that the knocked out trx2Δ could not survive the peroxide exposure. Although the endogenous thioredoxin’s expression allows surviving the peroxide treatment, the viability, measured as the percentage of live cells, was compromised. Similarly, expression of trx0 and D26I/A39K-trx0 complemented the lack of endogenous thioredoxin while A39K-trx0 was not able to complement (Fig 4b). Taken together, these experiments suggested that the engineered active site center of the thioredoxin could catalyze the reduction of disulfides bonds *in vivo.*

## DISCUSSION

Our results illustrate that optimizing the protein’s core hydrophobicity combined with surface charge distribution’s rearrangement results in an extensive stability enhancement while maintaining the functional properties of the thioredoxin. We showed that, for a previously designed hyper stable thioredoxin^29^, mutating a buried ionizable group results in a dramatic decrease in the redox capacity and, at the same time, increases protein stability by ten degrees. Combining the experimental screening of a library of mutations at selected positions with partial-least-square reconstruction of the whole mutational space, we found out that a sole alanine-to-lysine mutation in a solvent-exposed position is enough to restore the functional properties of the buried charge. Further, we found a very weak stability and redox activity trade-off that allows thioredoxin’s multi-feature optimization. Thus, we engineered thioredoxin’s active site to efficiently catalyze the reduction of disulfide bonds *in vitro* and stabilize the protein scaffold ten degrees. Lastly, we showed that the new thioredoxin could also function *in vivo.*

Protein engineering studies often seek improved stabilities. Several evolutionary-based strategies, such as back-to-consensus mutations^55–57^ or ancestral protein reconstruction^29,58,59^, have been well used to achieve protein stabilization. Also, directed evolution techniques have shown great promise in stabilizing proteins9,60. Overall, these methods have proven successful in stabilizing natural proteins. However, engineered proteins with the highest stabilities mostly come from *de novo* design^15,61–64^. Some of them display exceptionally high thermal stabilities; for example, Top7 and DRNN have melting temperatures of 133°C62,63 and 142°C15, respectively. One of the reasons for such impressive stabilities could be explained by the exclusively hydrophobic core of the designs^10^. However, natural proteins often display functional ionizable residues in their hydrophobic cores28,65 achieving functionality at the expense of thermodynamic stability^17,24,25^. Redesigning active sites with more stable configuration without compromising function is, thus, a challenge. Here, we have shown for the thioredoxin that redesigning its surface charge distribution nearby the active site can simulate the selected catalytic properties of a conserved and buried ionizable group. This approach allowed us to design one of the enzymes with the highest thermal stability ever reported. In fact, out of more than 32,000 entries in the 2021 version of the ProTherm database (https://web.iitm.ac.in/bioinfo2/prothermdb)^32^, only four natural proteins have similar or higher reported thermal stability. Overall, this approach optimizes the thioredoxin’s hydrophobic core while preserving its enzymatic activity and, therefore, opens a new strategy to stabilize enzymes.

We also showed that the redesigned thioredoxin can function *in vivo* yet does not fully compensate for the lack of the endogenous thioredoxin. This could be due to the multifunctional nature of the thioredoxin^49^. It reduces oxidized cysteine residues of several biological substrates, such as hydrogen peroxide, transcription factors^66^, unfolded proteins^67^, and kinases^68^ among others. Thus, thioredoxin is essential to keep intracellular protein disulfides reduced, prevent oxidative stress, regulate transcription factors’ activity, and assist other proteins’ folding. The active site center of the thioredoxin is plausibly the outcome of satisfying many of these functions. Here, we demonstrated that an alanine-to-lysine mutation in a solvent-exposed position is enough to restore the properties of the buried Asp_26_ to catalyze the reduction of disulfide bridges. However, we did not consider the role of the Asp_26_ in other functionalities and specific interactions with other proteins. Nevertheless, *in vivo* assays showed a slightly better complementation of D26I/A39K-trx0 relative to trx0, suggesting the nonessentiality of the Asp_26_. Another reason to explain the failure to fully complement the endogenous thioredoxin could be that the endogenous thioredoxin has coevolved with the organism’s protein network. However, the D26I/A39K-trx0 has not.

The design of new active sites from scratch is a grand challenge. For this purpose, a general approach implies burying, totally or partially, ionizable groups in the hydrophobic protein core^5,20,21,69^ because such buried groups often have perturbed physicochemical properties essential for their catalytic roles28,65. However, placing a charge in a hydrophobic pocket occurs at a thermodynamic cost^17,24,25^. As a result, a significant limitation to design enzymes with tailored functionalities resides in the thermostability of the protein scaffold^8^. Here, we have shown for the thioredoxin that modulating the surface charge distribution nearby its active site can simulate the catalytic properties of a buried charged group. Thus, our results open a new strategy that might help design new enzymes.

In summary, this work suggests to introduce functional innovation in a protein scaffold targeting those structural features more plastic. With this approach, we redesigned the thioredoxin active site center against conservation, expanding the evolutionary mutational space’s boundaries. We think this strategy can be generalized and exploited to stabilize natural proteins.

## METHODS

### Sequence alignment and conservation analysis

We have carried out a BLAST (https://www.ebi.ac.uk/Tools/sss/ncbiblast/) search using as query the E. *coli* thioredoxin sequence (PDB:2trx). The Swissprot database of November 2020 and the default options of the search were used. The retrieved sequences were aligned using the Smith-Waterman algorithm and, those sharing a minimum of 30% of identity and with entry names corresponding to thioredoxins (a total of 120), were retained. The consensus logo was generated with Weblogo (https://weblogo.berkeley.edu/logo.cgi).

### Gene library construction, and protein expression and purification

The combinatorial library was constructed by gene assembly mutagenesis^43^ and cloned into the plasmid pQE-80L with a 6xHis tag at the N-terminal to facilitate protein purification. The thioredoxin variants were expressed and purified as previously described^70^. Purity of the protein fractions were assessed by SDS-PAGE.

### *In vitro* activity assay

*In vitro* redox activity of the thioredoxin variants was determined by a turbidimetric assay using insulin as a substrate^41^. Briefly, we prepared protein solutions at pH 6.5 in phosphate buffer 0.1 M, 2 mM EDTA, and 0.5 mg/mL of insulin. The addition of DTT until 1 mM final concentration initiated the reaction that was monitored by measuring the absorbance at 650 nm over time. Then, the maximum value of the derivative of the kinetics was calculated. For each variant, 3 to 4 concentrations were assayed. The redox activity was quantified from the linear fit of the maximum derivative versus protein concentration (**Fig 1c** and **S1**).

### Partial least-squares analysis

PLS analyses were carried out with the program Unscrambler X from CAMO software using the NIPALS algorithm (**Fig 2**). The dependent variable was the logarithm value of the redox activity and was auto-scaled before the analysis. We used leave-one-out cross-validation. The number of latent variables retained was the Unscrambler program’s optimum value based on the mean square error of cross-validation.. The PLS analyses were carried with ten replica sets constructed from the original set through random resampling (bootstrapping). The number of latent variables retained depending on the replica set used. Typical values were much smaller than the numbers of dependent and independent variables involved. Illustrative plots of experimental versus predicted activities are given in **Fig 2c**.

### Thermostability experiments

Differential scanning calorimetry experiments were performed using an automatic VP-DSC microcalorimeter (VP-Capillary DSC (MicroCal, Malvern Panalytical)) with a 0.133 mL cell volume. Solutions of 0.5 mg/mL protein concentration in 50 mM HEPES at pH 7 were warmed up from 25°C until 130°C or 140°C at 210 K/min scan rate. Heat up to 140°C requires manipulating the calorimeter settings under the supervision of MicroCal. Unfortunately, the calorimeter that was manipulated to heat up until 140°C was damaged. Consequently, some experiments were performed up to 130°C. To prevent boiling above 100°C, overpressure was implemented. All thermograms were dynamically corrected using the software provided by MicroCal. The melting temperature (T_m_) was estimated at the maximum heat capacity of the calorimetric curve. For those calorimetry curves that do not show the pick of the scan in the studied range of temperatures, the T_m_ was inferred by fitting the data using a two-state model with fixed van’t Hoff enthalpy29.

### Principal component analysis

Principal component analysis was performed using R71, prior to the analysis both variables (Redox activity and Thermal stability) were autoscaled.

### Genome integration of thioredoxins in yeast

Genes encoding the ancestral thioredoxin variants trx0, trx0 D26I, trx0 D26I/A39K, and S. *cerevisiae* trx2 were subcloned downstream of the yeast GPD promoter into an M3925 plasmid (trp1::KanMX3) with a cassette for genome integration^72^. For genome integration, the cassette was amplified by PCR and transformed into BY4741 cells. Transformants were selected by G418 resistance, and correct integration was assessed by sequencing.

### Growth assays of yeast strains

Yeast strains were grown for 2 days in YPD containing 200μg/mL of G418 at 30°C until reaching the stationary phase. Then, strains were diluted 1:100 in YPD with concentrations of 0, 3, and 4 mM H_2_O_2_ in a 96-well plate for a final volume of 150μL per well. Three replicates of each strain were done for each condition. Strains were grown under constant shaking for 60h at 30°C in a plate reader. Absorbance at 600 nm was continuously monitored throughout the experiment. Analysis of the Growth curves (**Fig 4** and **Table S2**) were done with the growthrate R package^73^.

### Plate-based spot viability assay

Yeast cultures of 2 mL of YPD supplemented with G418 200 μg/mL were grown in a 96-well plate, at 30°C with saturated humidity until saturation. Then, from this preculture, two new cultures were prepared at 0.1 OD, the control and the oxidatively stressed culture. The control culture was prepared in YPD, and from this one, five more cultures were prepared by two times dilution. In total, for each yeast strain, six solutions in YPD with an absorbance at 600 nm of 0.1, 0.05, 0.025, 0.0125, 0.006 and 0.003 respectively. We spotted 5 μL of each culture in a YPD-agar plate, let it dry, and left it at 30°C for two days. Since absorbance 1 at 600 nm correspond around 0.5 *10^7^ cells/mL, we spotted 2500, 1250, 625, 312, 160, and 80 cells in each drop (**Fig 4b**). For the oxidative stress shock, we prepared six solutions in YPD supplemented with 4mM H_2_O_2_ with an absorbance at 600 nm of 0.1, 0.05, 0.025, 0.0125, 0.006, and 0.003. The cultures were incubated 24h at 30°C with saturated humidity. Then, we spotted 5 μL of each culture in a YPD-agar plate, let it dry, and left it at 30°C for two days (**Fig 4b**).

## AUTHOR CONTRIBUTION

M.L.R.R., B.I.M., and H.G.S. designed the experiments. M.L.R.R. and H.G.S. performed the experiments. M.L.R.R. and H.G.S. analyzed the data, M.L.R.R, H.G.S. and B.I.M. wrote the manuscript.

## ACKNOWLEDGEMENTS

We thank Dr. Jose Manuel Sanchez Ruiz for invaluable input throughout the realization of this work, Dr. Agnes Toth-Petrozcy for useful discussions on the manuscript, Dr. Emmanuel D. Levy for sharing the yeast strains and the plasmids with the cassettes for genome integration and Dr. Paola Laurino and Dr. Benjamin Clifton for feedback and comments that greatly improved the manuscript. This work was supported by European Regional Development Fund funds and grants (CSD2009-00088 and BIO2012-34937), by the Spanish Ministry of Economy and Competitiveness (grant BIO2015-66426-R), and by the “Junta de Andalucia” (grant P09-CVI-5073). M.L.R.R. and H.G.S. received support from the Koshland Foundation and a McDonald-Leapman Grant.

## SUPPLEMENTARY INFORMATION

**Table S1.** Values of δ_*ij*_ according with the type of reside placed in positions *i* and *j*. δ is a vector of values (1, 0, 0) when the position is occupied by a neutral residue, (0, 1, 0) when positively charged residue occupies it, and (0, 0, 1) when a negatively charged residue occupies it; δ_ij_ = δ_*i*_·δ_*j*_. takes different values depending on the type of residue in positions i and j.

| position $i$ | $\delta_i$ | position $j$ | $\delta_j$ | $\delta_{ij}$ |
| --- | --- | --- | --- | --- |
| neutral | (1,0,0) | neutral | (1,0,0) | (0,0,1,0,0,0,0,0,0,0,0,0,0,0,0) |
| neutral | (1,0,0) | positive | (0,1,0) | (0,0,0,1,0,0,0,0,0,0,0,0,0,0,0) |
| neutral | (1,0,0) | negative | (0,0,1) | (0,0,0,0,1,0,0,0,0,0,0,0,0,0,0) |
| positive | (0,1,0) | neutral | (1,0,0) | (0,0,0,0,0,1,0,0,0,0,0,0,0,0,0) |
| positive | (0,1,0) | positive | (0,1,0) | (0,0,0,0,0,0,1,0,0,0,0,0,0,0,0) |
| positive | (0,1,0) | negative | (0,0,1) | (0,0,0,0,0,0,0,1,0,0,0,0,0,0,0) |
| negative | (0,0,1) | neutral | (1,0,0) | (0,0,0,0,0,0,0,0,1,0,0,0,0,0,0) |
| negative | (0,0,1) | positive | (0,1,0) | (0,0,0,0,0,0,0,0,0,1,0,0,0,0,0) |
| negative | (0,0,1) | negative | (0,0,1) | (0,0,0,0,0,0,0,0,0,0,1,0,0,0,0) |

**Table S2.** Parameters obtained using fit_easylinear of Growthrate R package^73^. ODt_0_ is the OD value at the starting point. *μ*_max_ is the value of the maximum growth rate. The time that the culture takes to grow exponentially is the lag phase.

| strain | H<sub>2</sub>O<sub>2</sub> (mM) | OD t<sub>0</sub> | $\mu_{\max}$ (h<sup>-1</sup>) | lag phase (h) |
| --- | --- | --- | --- | --- |
| trx2Δ | 0 | 0.01 | 0.32 | 7.68 |
| trx2Δ | 3 | 0.00 | 0.30 | 25.19 |
| trx2Δ | 4 | 0.00 | 0.30 | 48.27 |
| A39K-D26I-trx0/trx2Δ | 0 | 0.00 | 0.30 | 8.11 |
| A39K-D26I-trx0/trx2Δ | 3 | 0.00 | 0.28 | 21.45 |
| A39K-D26I-trx0/trx2Δ | 4 | 0.00 | 0.26 | 40.57 |
| trx0/trx2Δ | 0 | 0.01 | 0.31 | 8.33 |
| trx0/trx2Δ | 3 | 0.00 | 0.29 | 22.90 |
| trx0/trx2Δ | 4 | 0.00 | 0.29 | 43.32 |
| D26I-trx0/trx2Δ | 0 | 0.01 | 0.31 | 8.64 |
| D26I-trx0/trx2Δ | 3 | 0.00 | 0.29 | 24.69 |
| D26I-trx0/trx2Δ | 4 | 0.00 | 0.27 | 49.73 |
| trx2/trx2Δ | 0 | 0.01 | 0.25 | 7.49 |
| trx2/trx2Δ | 3 | 0.00 | 0.30 | 13.40 |
| trx2/trx2Δ | 4 | 0.00 | 0.29 | 19.09 |

**Figure S1.**
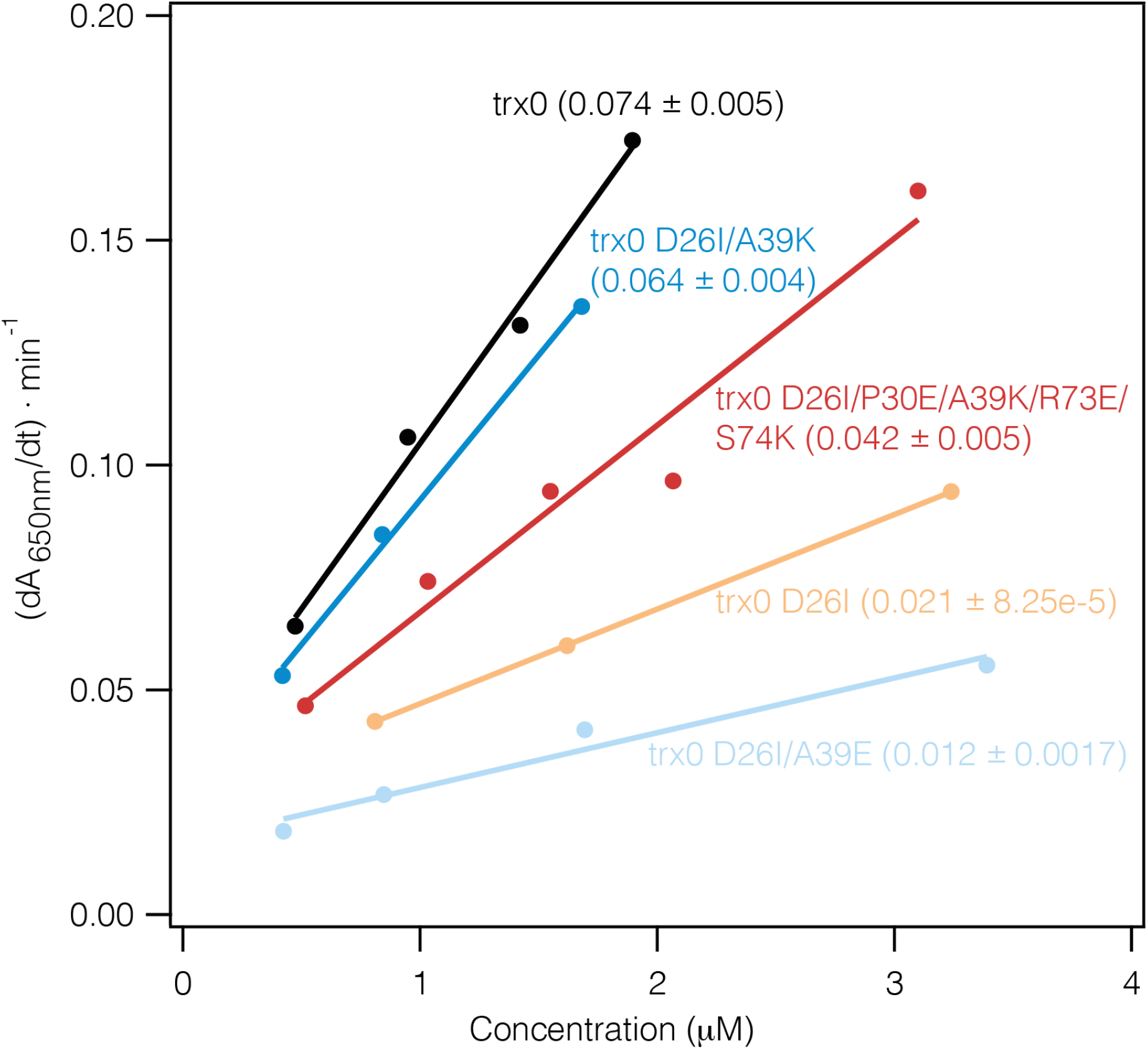
Reductase activity determination of several representative thioredoxin variants. We determine the activity of each variant from the slope of the linear relationship between the maximum value of the 650 nm absorbance resulting from the insulin reduction curves versus protein concentration. In the plot, we show several representative variants, including trx0, the background of the library trx0 D26I, and trx0 D26I A39K.

**Figure S2.**
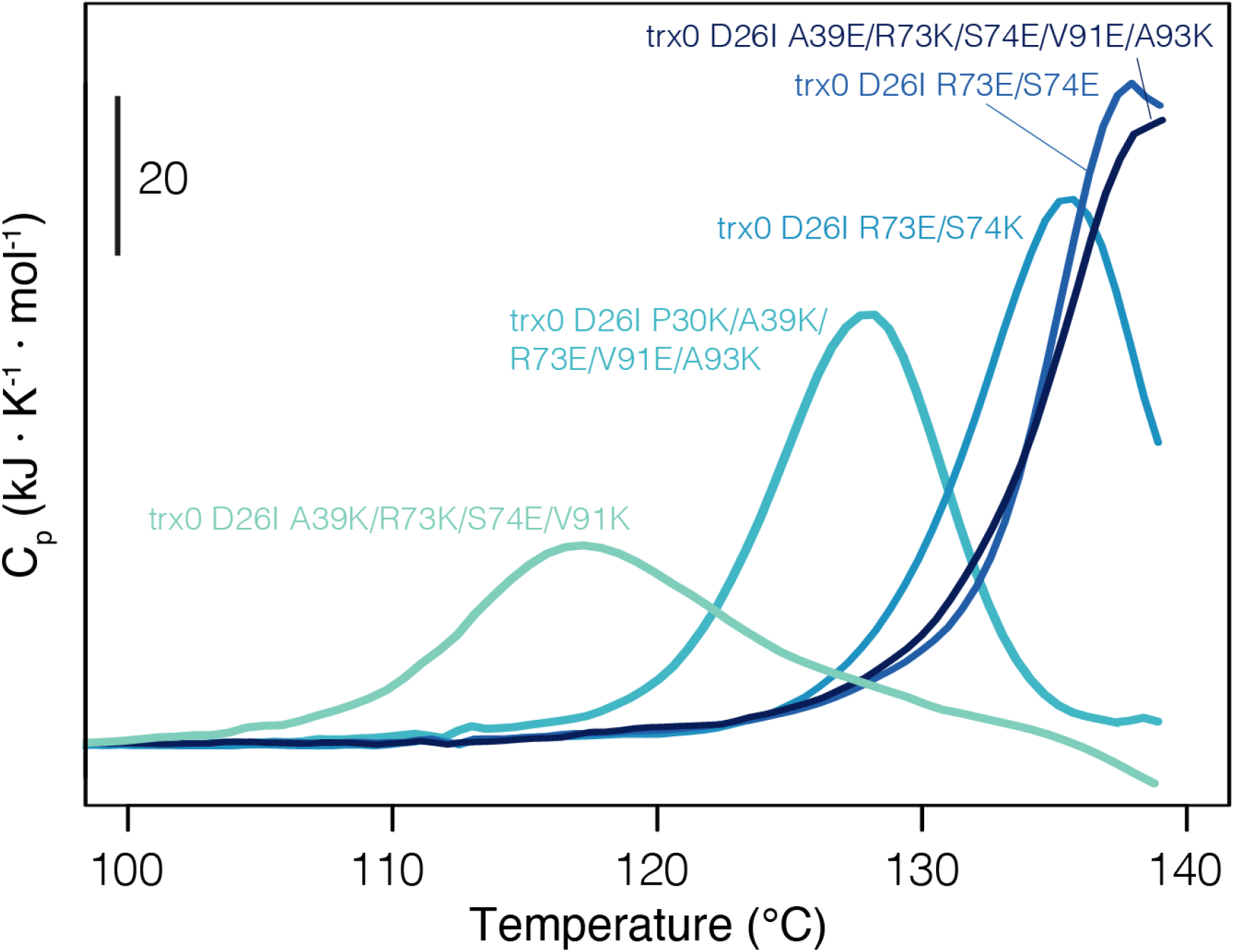
Differential scanning calorimetry experiments of several representative thioredoxin variants. Thioredoxin samples of 0.5 mg/mL protein concentration in 50 mM HEPES at pH7 were heated from 25°C to 140°C at 210 K/min scan rate. Melting temperatures were attributed to the maximum heat capacity of the calorimetric curves. In the plot, we show several representative thioredoxin library variants.

**Figure S3.**
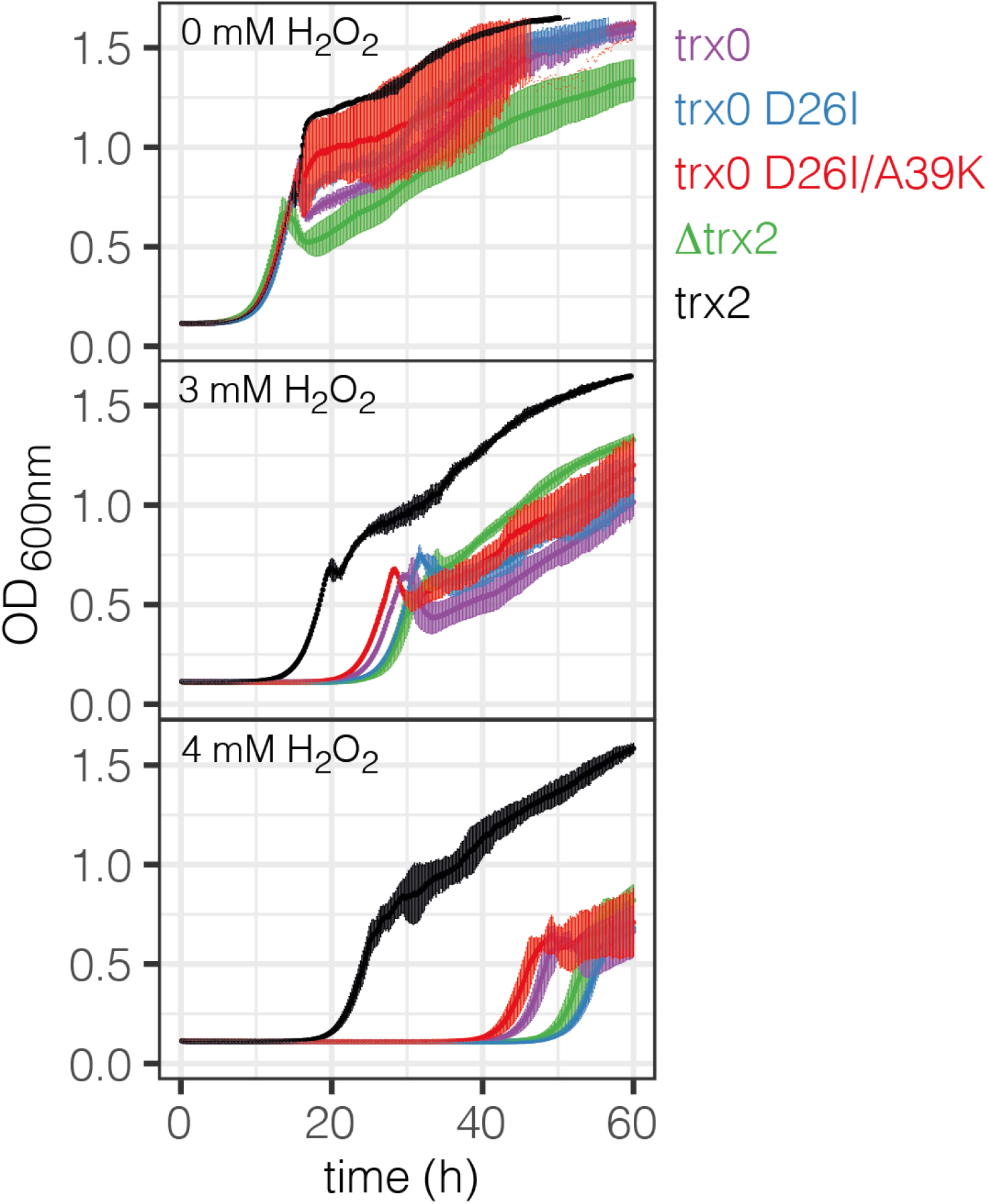
Growth curves of yeast strains at different H_2_O_2_ concentrations. The yeast strain lacking the trx2 gene (Δtrx2) and the strains where trx0, trx0 D26I, trx0 D26I/A39K, and trx2 were genome-integrated on the Δtrx2 background were grown at 30°C in YPD in the presence of 0, 3 and 4 mM H_2_O_2_. Three replicates of each strain were done for each condition. Strains were grown under constant shaking. Absorbance at 600nm was continuously monitored.

## REFERENCES

1. Magliery, T. J. Protein stability: computation, sequence statistics, and new experimental methods. Curr. Opin. Struct. Biol. 33, 161–168 (2015).

2. Tokuriki, N., Stricher, F., Serrano, L. & Tawfik, D. S. How protein stability and new functions trade off. PLoS Comput. Biol. 4, e1000002 (2008).

3. Khersonsky, O. et al. Bridging the gaps in design methodologies by evolutionary optimization of the stability and proficiency of designed Kemp eliminase KE59. Proc. Natl. Acad. Sci. U. S. A. 109, 10358–10363 (2012).

4. Bloom, J. D., Labthavikul, S. T., Otey, C. R. & Arnold, F. H. Protein stability promotes evolvability. Proc. Natl. Acad. Sci. U. S. A. 103, 5869–5874 (2006).

5. Röthlisberger, D. et al. Kemp elimination catalysts by computational enzyme design. Nature 453, 190–195 (2008).

6. Fleishman, S. J. et al. Computational design of proteins targeting the conserved stem region of influenza hemagglutinin. Science 332, 816–821 (2011).

7. Wijma, H. J., Floor, R. J. & Janssen, D. B. Structure- and sequence-analysis inspired engineering of proteins for enhanced thermostability. Current Opinion in Structural Biology vol. 23 588–594 (2013).

8. Goldenzweig, A. & Fleishman, S. J. Principles of Protein Stability and Their Application in Computational Design. Annu. Rev. Biochem. 87, 105–129 (2018).

9. Chandler, P. G. et al. Strategies for Increasing Protein Stability. Methods Mol. Biol. 2073, 163–181 (2020).

10. Baker, D. What has de novo protein design taught us about protein folding and biophysics? Protein Sci. 28, 678–683 (2019).

11. Munson, M. et al. What makes a protein a protein? Hydrophobic core designs that specify stability and structural properties. Protein Sci. 5, 1584–1593 (1996).

12. Malakauskas, S. M. & Mayo, S. L. Design, structure and stability of a hyperthermophilic protein variant. Nat. Struct. Biol. 5, 470–475 (1998).

13. Bolon, D. N. & Mayo, S. L. Polar residues in the protein core of Escherichia coli thioredoxin are important for fold specificity. Biochemistry 40, 10047–10053 (2001).

14. Borgo, B. & Havranek, J. J. Automated selection of stabilizing mutations in designed and natural proteins. Proc. Natl. Acad. Sci. U. S. A. 109, 1494–1499 (2012).

15. Murphy, G. S. et al. Increasing sequence diversity with flexible backbone protein design: the complete redesign of a protein hydrophobic core. Structure 20, 1086–1096 (2012).

16. Rocklin, G. J. et al. Global analysis of protein folding using massively parallel design, synthesis, and testing. Science 357, 168–175 (2017).

17. Isom, D. G., Castañeda, C. A., Cannon, B. R., Velu, P. D. & García-Moreno E, B. Charges in the hydrophobic interior of proteins. Proc. Natl. Acad. Sci. U. S. A. 107, 16096–16100 (2010).

18. Liu, J., Swails, J., Zhang, J. Z. H., He, X. & Roitberg, A. E. A Coupled Ionization-Conformational Equilibrium Is Required To Understand the Properties of Ionizable Residues in the Hydrophobic Interior of Staphylococcal Nuclease. J. Am. Chem. Soc. 140, 1639–1648 (2018).

19. Narayan, A. & Naganathan, A. N. Switching Protein Conformational Substates by Protonation and Mutation. J. Phys. Chem. B 122, 11039–11047 (2018).

20. Risso, V. A. et al. De novo active sites for resurrected Precambrian enzymes. Nat. Commun. 8, 16113 (2017).

21. Korendovych, I. V. et al. Design of a switchable eliminase. Proc. Natl. Acad. Sci. U. S. A. 108, 6823–6827 (2011).

22. Jiang, L. et al. De novo computational design of retro-aldol enzymes. Science 319, 1387–1391 (2008).

23. Althoff, E. A. et al. Robust design and optimization of retroaldol enzymes. Protein Sci. 21, 717–726 (2012).

24. Garcia-Seisdedos, H., Ibarra-Molero, B. & Sanchez-Ruiz, J. M. How many ionizable groups can sit on a protein hydrophobic core? Proteins: Struct. Funct. Bioinf. 80, 1–7 (2012).

25. Pace, C. N., Grimsley, G. R. & Scholtz, J. M. Protein ionizable groups: pK values and their contribution to protein stability and solubility. J. Biol. Chem. 284, 13285–13289 (2009).

26. Kougentakis, C. M. et al. Anomalous Properties of Lys Residues Buried in the Hydrophobic Interior of a Protein Revealed with 15N-Detect NMR Spectroscopy. J. Phys. Chem. Lett. 9, 383–387 (2018).

27. Tóth-Petróczy, Á. & Tawfik, D. S. Slow protein evolutionary rates are dictated by surface–core association. Proceedings of the National (2011).

28. Pey, A. L., Rodriguez-Larrea, D., Gavira, J. A., Garcia-Moreno, B. & Sanchez-Ruiz, J. M. Modulation of buried ionizable groups in proteins with engineered surface charge. J. Am. Chem. Soc. 132, 1218–1219 (2010).

29. Romero-Romero, M. L., Risso, V. A., Martinez-Rodriguez, S., Ibarra-Molero, B. & Sanchez-Ruiz, J. M. Engineering ancestral protein hyperstability. Biochem. J 473, 3611–3620 (2016).

30. Holmgren, A., Söderberg, B. O., Eklund, H. & Brändén, C. I. Three-dimensional structure of Escherichia coli thioredoxin-S2 to 2.8 A resolution. Proc. Natl. Acad. Sci. U. S. A. 72, 2305–2309 (1975).

31. Gleason, F. K. Mutation of conserved residues in Escherichia coli thioredoxin: effects on stability and function. Protein Sci. 1, 609–616 (1992).

32. Nikam, R., Kulandaisamy, A., Harini, K., Sharma, D. & Gromiha, M. M. ProThermDB: thermodynamic database for proteins and mutants revisited after 15 years. Nucleic Acids Res. 49, D420–D424 (2021).

33. Holmgren, A. Thioredoxin. Annu. Rev. Biochem. 54, 237–271 (1985).

34. Holmgren, A. Thioredoxin structure and mechanism: conformational changes on oxidation of the active-site sulfhydryls to a disulfide. Structure 3, 239–243 (1995).

35. LeMaster, D. M., Springer, P. A. & Unkefer, C. J. The Role of the Buried Aspartate of Escherichia coli Thioredoxin in the Activation of the Mixed Disulfide Intermediate. Journal of Biological Chemistry vol. 272 29998–30001 (1997).

36. Menchise, V. et al. Crystal structure of the wild-type and D30A mutant thioredoxin h of Chlamydomonas reinhardtii and implications for the catalytic mechanism. Biochem. J 359, 65–75 (2001).

37. Chivers, P. T. & Raines, R. T. General acid/base catalysis in the active site of Escherichia coli thioredoxin. Biochemistry 36, 15810–15816 (1997).

38. Perez-Jimenez, R. et al. Single-molecule paleoenzymology probes the chemistry of resurrected enzymes. Nat. Struct. Mol. Biol. 18, 592–596 (2011).

39. Ingles-Prieto, A. et al. Conservation of protein structure over four billion years. Structure 21, 1690–1697 (2013).

40. Eklund, H., Gleason, F. K. & Holmgren, A. Structural and functional relations among thioredoxins of different species. Proteins 11, 13–28 (1991).

41. Holmgren, A. Thioredoxin catalyzes the reduction of insulin disulfides by dithiothreitol and dihydrolipoamide. J. Biol. Chem. 254, 9627–9632 (1979).

42. Fox, R. J. et al. Improving catalytic function by ProSAR-driven enzyme evolution. Nat. Biotechnol. 25, 338–344 (2007).

43. Garcia-Seisdedos, H., Ibarra-Molero, B. & Sanchez-Ruiz, J. M. Probing the mutational interplay between primary and promiscuous protein functions: a computational-experimental approach. PLoS Comput. Biol. 8, e1002558 (2012).

44. Berland, M., Offmann, B., André, I., Remaud-Siméon, M. & Charton, P. A web-based tool for rational screening of mutants libraries using ProSAR. Protein Eng. Des. Sel. 27, 375–381 (2014).

45. Partial Least Squares Regression. http://methods.sagepub.com/reference/the-sage-encyclopedia-of-social-science-research-methods/n690.xml doi:10.4135/9781412950589.n690.

46. Lever, J., Krzywinski, M. & Altman, N. Principal component analysis. Nature Methods vol. 14 641–642 (2017).

47. Giaever, G. et al. Functional profiling of the Saccharomyces cerevisiae genome. Nature 418, 387–391 (2002).

48. Kuge, S. & Jones, N. YAP1 dependent activation of TRX2 is essential for the response of Saccharomyces cerevisiae to oxidative stress by hydroperoxides. EMBO J. 13, 655–664 (1994).

49. Arnér, E. S. & Holmgren, A. Physiological functions of thioredoxin and thioredoxin reductase. Eur. J. Biochem. 267, 6102–6109 (2000).

50. Kumar, J. K., Tabor, S. & Richardson, C. C. Proteomic analysis of thioredoxin-targeted proteins in Escherichia coli. Proc. Natl. Acad. Sci. U. S. A. 101, 3759–3764 (2004).

51. Arts, I. S., Vertommen, D., Baldin, F., Laloux, G. & Collet, J.-F. Comprehensively Characterizing the Thioredoxin Interactome In Vivo Highlights the Central Role Played by This Ubiquitous Oxidoreductase in Redox Control. Mol. Cell. Proteomics 15, 2125–2140 (2016).

52. Oughtred, R. et al. The BioGRID database: A comprehensive biomedical resource of curated protein, genetic, and chemical interactions. Protein Sci. 30, 187–200 (2021).

53. Tran, K. & Green, E. M. Assessing Yeast Cell Survival Following Hydrogen Peroxide Exposure. Bio Protoc 9, (2019).

54. Kwolek-Mirek, M. & Zadrag-Tecza, R. Comparison of methods used for assessing the viability and vitality of yeast cells. FEMS Yeast Res. 14, 1068–1079 (2014).

55. Oda, K. et al. Consensus mutagenesis approach improves the thermal stability of system xc - transporter, xCT, and enables cryo-EM analyses. Protein Sci. 29, 2398–2407 (2020).

56. Georgoulis, A. et al. Consensus protein engineering on the thermostable histone-like bacterial protein HUs significantly improves stability and DNA binding affinity. Extremophiles 24, 293–306 (2020).

57. Yao, H., Cai, H. & Li, D. Thermostabilization of Membrane Proteins by Consensus Mutation: A Case Study for a Fungal ?8-7 Sterol Isomerase. J. Mol. Biol. 432, 5162–5183 (2020).

58. Risso, V. A., Gavira, J. A., Mejia-Carmona, D. F., Gaucher, E. A. & Sanchez-Ruiz, J. M. Hyperstability and substrate promiscuity in laboratory resurrections of Precambrian ?-lactamases. J. Am. Chem. Soc. 135, 2899–2902 (2013).

59. Risso, V. A., Sanchez-Ruiz, J. M. & Ozkan, S. B. Biotechnological and protein-engineering implications of ancestral protein resurrection. Curr. Opin. Struct. Biol. 51, 106–115 (2018).

60. Höcker, B. Directed evolution of (betaalpha)(8)-barrel enzymes. Biomol. Eng. 22, 31–38 (2005).

61. Lin, Y.-R. et al. Control over overall shape and size in de novo designed proteins. Proc. Natl. Acad. Sci. U. S. A. 112, E5478–85 (2015).

62. Koga, R. et al. Robust folding of a de novo designed ideal protein even with most of the core mutated to valine. Proc. Natl. Acad. Sci. U. S. A. 117, 31149–31156 (2020).

63. Kuhlman, B. et al. Design of a novel globular protein fold with atomic-level accuracy. Science 302, 1364–1368 (2003).

64. Marcos, E. et al. De novo design of a non-local ?-sheet protein with high stability and accuracy. Nat. Struct. Mol. Biol. 25, 1028–1034 (2018).

65. Isom, D. G., Castañeda, C. A., Cannon, B. R. & García-Moreno, B. Large shifts in pKa values of lysine residues buried inside a protein. Proc. Natl. Acad. Sci. U. S. A. 108, 5260–5265 (2011).

66. Matthews, J. R., Wakasugi, N., Virelizier, J.-L., Yodoi, J. & Hay, R. T. Thiordoxin regulates the DNA binding activity of NF-χB by reduction of a disulphid bond involving cysteine 62. Nucleic Acids Research vol. 20 3821–3830 (1992).

67. Kern, R., Malki, A., Holmgren, A. & Richarme, G. Chaperone properties of Escherichia coli thioredoxin and thioredoxin reductase. Biochem. J 371, 965–972 (2003).

68. Saitoh, M. et al. Mammalian thioredoxin is a direct inhibitor of apoptosis signal-regulating kinase (ASK) 1. EMBO J. 17, 2596–2606 (1998).

69. Burton, A. J., Thomson, A. R., Dawson, W. M., Brady, R. L. & Woolfson, D. N. Installing hydrolytic activity into a completely de novo protein framework. Nat. Chem. 8, 837–844 (2016).

70. Romero-Romero, M. L., Inglés-Prieto, A., Ibarra-Molero, B. & Sanchez-Ruiz, J. M. Highly anomalous energetics of protein cold denaturation linked to folding-unfolding kinetics. PLoS One 6, e23050 (2011).

71. Team, R. C. RA language and environment for statistical computing, Vienna, Austria; 2019. Available in: http://www.R-project.org/Cited 5, (2019).

72. Garcia-Seisdedos, H., Levin, T., Shapira, G. & Freud, S. Mutants libraries reveal negative design shielding proteins from mis-assembly and re-localization in cells. bioRxiv (2021).

73. Hall, B. G., Acar, H., Nandipati, A. & Barlow, M. Growth rates made easy. Mol. Biol. Evol. 31, 232–238 (2014).

